# A Mac2-positive progenitor-like microglial population survives independent of CSF1R signaling in adult mouse brain

**DOI:** 10.1101/722090

**Authors:** Lihong Zhan, Peter Dongmin Sohn, Yungui Zhou, Yaqiao Li, Li Gan

**Affiliations:** Gladstone Institute of Neurological Diseases, San Francisco, CA 94158, USA; Department of Neurology, University of California, San Francisco, CA 94158, USA; Helen and Robert Appel Alzheimer’s Disease Institute, Brain and Mind Research Institute, Weill Cornell Medicine, New York, NY10065, USA

**Keywords:** Microglia, Myeloid Cells, CSF1R signaling, Microglia Homeostasis, Single-cell RNA Sequencing, Mac2, Adult Microglial Progenitors

## Abstract

Microglia are the resident myeloid cells in the central nervous system (CNS). The majority of microglial population relies on Csf1r signaling for survival and maintenance. However, a small subset of microglia in the murine brain can survive without Csf1r signaling, and reestablishes homeostasis after Csf1r signaling returns. Using single-cell RNA-seq, we characterized the heterogeneous microglial populations under Csf1r inhibition, including microglia lacking homeostatic markers and populations with elevated markers of monocytes, granulocytes and dendritic cells. Importantly, Mac2 is distinctively expressed in a subset of Csf1r-independent microglia cells, which were highly proliferative and shared striking similarities with those of microglial progenitors in yolk sac and early embryos. Lineage-tracing revealed that the Mac2+ population is of microglial origin and does not come from periphery monocytes. In non-treated mouse brains, Mac2+ microglia exhibited progenitor transcriptomic signature indistinguishable from those survived csf1r inhibition, supporting Mac2+ progenitor-like cells are present among homeostatic microglia.

## INTRODUCTION

Microglia are the primary innate immune cells in the CNS, capable of mounting inflammatory responses and phagocytosis. They can be distinguished from other CNS cell types by the distinctive ramified morphology and expression of common myeloid markers including Cd11b and ionized calcium binding adaptor molecule 1 (Iba1). In addition to immune functions, microglia carry out a multitude of neurotrophic functions during CNS development and homeostasis (Kierdorf and Prinz, 2017). Microglia also play critical pathological roles in a wide spectrum of neurodegenerative conditions, including Alzheimer’s disease, Parkinson’s disease, Huntington’s disease, and Amyotrophic lateral sclerosis (Hickman et al., 2018). A number of disease genes were found to be highly expressed in microglia (Hansen et al., 2018), such as Trem2 (Abduljaleel et al., 2014; Jonsson et al., 2013) and Progranulin (Baker et al., 2006), highlighting the importance of microglia in neurodegenerative diseases.

Unlike other CNS glial cells, microglia originate from the embryonic mesoderm and follow a convoluted developmental journey (Rezaie and Male, 2002). It starts with the emergence of c-kit+ erythromyeloid progenitors in the yolk sac, known as primitive hematopoiesis, which then influx into the developing parenchyma via circulation (Ginhoux et al., 2010) in an Irf-8, Pu.1-dependent manner (Kierdorf et al., 2013). Seeded microglial progenitors persist in the CNS and continue to expand and mature until adulthood (Matcovitch-Natan et al., 2016). In general, developing microglia can be distinguished by well-defined developmental intervals from the yolk sac to the adult, and transcriptional programs associated with each of these stages have been meticulously mapped (Matcovitch-Natan et al., 2016). In particular, homeostatic maturation in microglia requires the transcriptional factor Mafb (Matcovitch-Natan et al., 2016) as well as TGF-beta signaling (Butovsky et al., 2014; Zoller et al., 2018), and can be distinguished by homeostatic markers such as Tmem119 (Bennett et al., 2016) and P2ry12 (Haynes et al., 2006). Interestingly, we recently discovered that adult newborn microglia follow a similar maturation path (Zhan et al., 2019), suggesting that the developmental plasticity of microglia in the adult brain might be an underlying feature of microglial homeostasis (Santambrogio et al., 2001). Unlike other tissue myeloid populations such as monocytes and macrophages, the resident microglial pool receives no significant replenishment from circulation and is internally maintained by self-renewal (Ajami et al., 2007; Mildner et al., 2007), even under conditions of acute ablation (Bruttger et al., 2015; Huang et al., 2018; Zhan et al., 2019). It is thus not surprising that microglia have an extremely long half-life (Lawson et al., 1992), most recently estimated to be 7.5-15 months in the murine CNS (Fuger et al., 2017; Tay et al., 2017; Zhan et al., 2019). In contrast, other myeloid populations such as classical monocytes have a half-life of less than 24 hours (van Furth and Cohn, 1968; Yona et al., 2013), and require constant replenishment from a Cx3cr1-population in the bone marrow (Fogg et al., 2006).

Csf1r signaling is critical for microglial survival and maintenance. Loss-of-function mutation is either of its two natural ligands: Csf1 and IL-34, results in significant reduction of microglia count (Greter et al., 2012; Wegiel et al., 1998). Null mutations in *Csf1r* remove 99.7% microglia, but a few morphologically distinctive microglia near the hippocampus and piriform cortex remain (Erblich et al., 2011). In addition, the Csf1r inhibitor PLX5622 (PLX) has been widely used as a research tool to acutely remove microglia. While depletion efficiency varies, complete microglial ablation has never been reported (Acharya et al., 2016; Huang et al., 2018; Rice et al., 2017; Zhan et al., 2019). These studies suggest that the adult microglial pool includes a population that does not require Csf1r signaling for survival. Remarkably, the microglial pool can be rapidly regenerated after termination of PLX administration. While an earlier study proposed a hidden nestin+ progenitor pool responsible for repopulation (Elmore et al., 2014), we and others found that the remaining microglia were solely responsible for microglial repopulation (Huang et al., 2018; Zhan et al., 2019), consistent with the notion that a subpopulation of microglia exhibit progenitor-like features. Our current study applied single-cell RNA-sequencing (scRNA-seq) to examine the resilient microglial population after acute Csf1r inhibition with PLX. Among the PLX-resistant microglia were a subpopulation of Mac2+ cells, which were also discovered in the homeostatic microglial pool yet displayed unexpected progenitor signatures. We also examined the role of the Mac2+ microglia during adult microgliogenesis in a microglia depletion/repopulation model.

## RESULTS

### Single cell RNA-seq profiling of microglia under Csf1r inhibition and early repopulation

To examine the transcriptome profiles of the Csf1r inhibitor-resistant microglial population, we performed scRNA-seq on the remaining microglia from C57/BL6J mice that were treated with PLX diet (D0) (Fig. 1A). To investigate early stage adult newborn microglia, we also included mice that were switched to a control diet for 2 days after PLX treatment (D2) (Fig 1A). Microglia from age-matched non-treated mice were used as control (Ctrl) (Fig 1A). Similar to our previous results (Zhan et al., 2019), oral dosing of 1200 mg/kg PLX in C57/BL6J mice for two weeks resulted 92.93 – 96.2% removal of CD11b+ myeloid cells in the CNS (Fig S1). Single cell suspensions were stained with CD11b antibody for myeloid population purification via fluorescence activated cell sorting (FACS) (Fig S1), followed by the 10x genomics single cell RNA-seq platform (Fig 1B).

**Fig 1.**
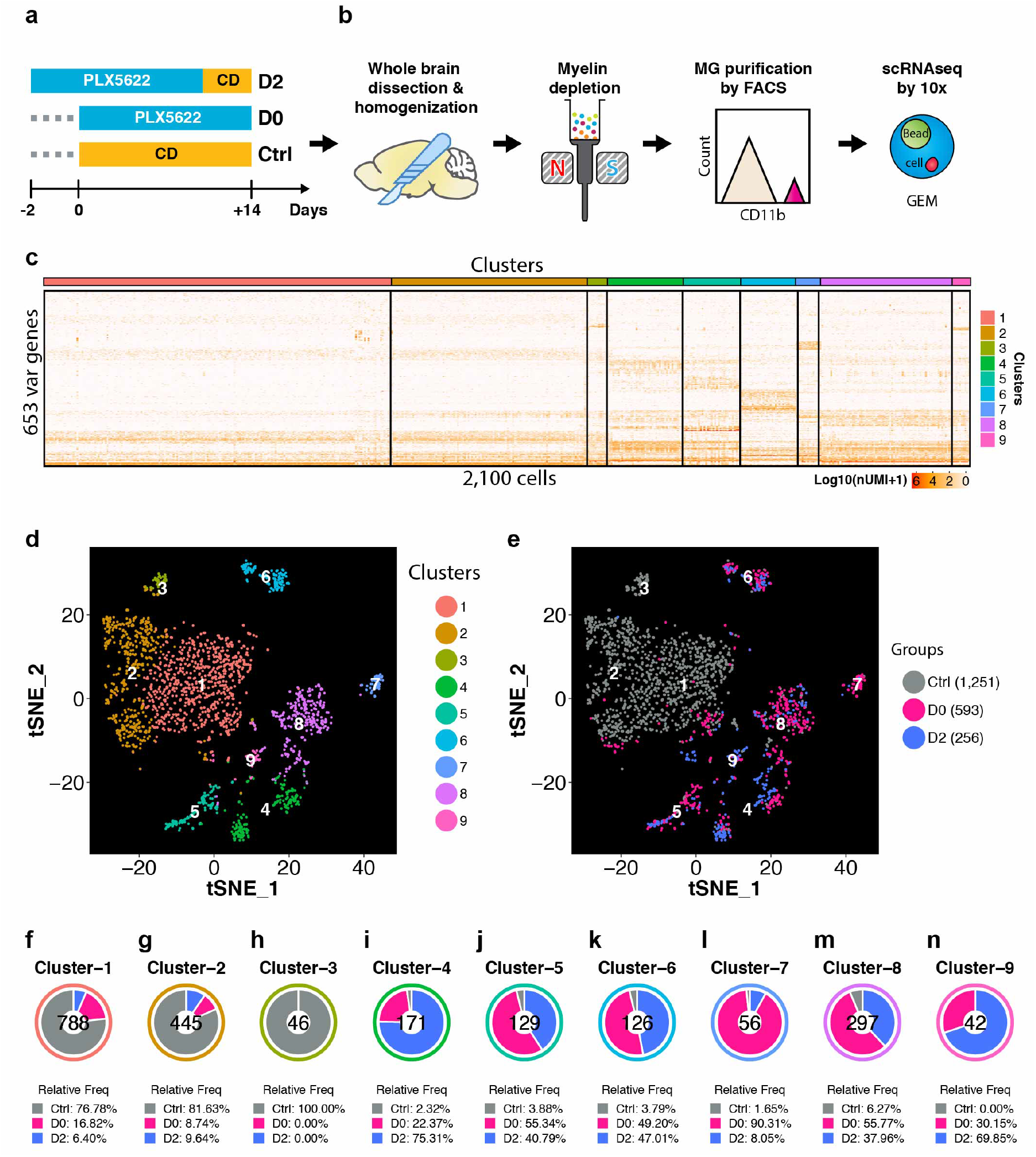
Single-cell RNA-seq profiling of microglia under Csf1r inhibition and early repopulation. (**a**) Experimental design for microglia depletion and repopulation. Mice were placed on PLX diet (1200 mg/kg) for 14 days to deplete microglia (D0). The early stage microglial repopulation (D2) group was switched to control diet (CD) for 2 days. Microglia from each mouse were collected in the same day. A total of 4 female C57/BL6J mice (5 Mo) were used: Ctrl (n=1); D0 (n=2, pooled together for FACS); D2 (n=1); (**b**) Workflow of adult microglia isolation procedures for scRNA-seq capture. Detailed description in methods. (**c**) Heatmap showing the most variable genes detected from 2,100 cells after initial data filtering. Log-transformed total UMI was plotted. (**d**) tSNE plot showing 9 distinctive clusters identified from the scRNA-seq data. (**e**) tSNE plot showing distribution of cells from each treatment group in 9 clusters. Number of cells from each group are: Ctrl (1251 cells); D0 (593 cells); D2 (256 cells). (**f-n**) Donut charts showing relative frequency (freq) of cells from three treatment groups distributed in each cluster. Relative cell frequency was calculated by normalizing to the total number of cells captured in each treatment group. The number of cells found in each cluster is shown at the center.

After data filtering with QC metrics (Fig. S2), we obtained a total of 2,100 cells at a sequencing depth of about 48,000 reads/cell (Fig 1C). A total of 653 variable genes were detected and the most variable genes were detected from 2100 cells after initial data filtering (Fig 1C). Although the number of cells sequenced from Ctrl, D0 and D2 differed, the sequencing quality was comparable, as evidenced by UMI, total number of genes, and percentage of mitochondrial genes per cell (Fig. S2). Following initial principle component analyses (Fig. S2), further data dimensional reduction using t-distributed stochastic neighbor embedding (tSNE) (van der Maaten, 2008) identified 9 distinctive cell clusters (Fig 1D). Of note, Clusters 1-3 consisted primarily of the naive microglia from control mice (Fig 1E), whereas cells from microglial depletion group (D0) and early repopulation group (D2) were grouped together in Clusters 4-9 (Fig 1E). Specifically, in Clusters 1–3, 76%–100% of microglia were from control mice (Fig. 1F–H) whereas less than 7% of microglia were from control mice in Clusters 4–9 (Fig. 1I–N). The overlapping representation of D0 and D2 microglia in Clusters 4–9 suggests that the early stage repopulating microglia are similar to those under Csf1r inhibition.

### Microglial homeostatic signatures are lost in remaining microglia under PLX treatment or early stage repopulation

In stark contrast to clusters derived from steady-state cells from control mice (Clusters 1-3), microglial homeostatic genes were significantly down-regulated in the clusters enriched with D0 and D2 cells (Cluster 4-9) (Fig 2A), including signature genes such as Tmem119 (Fig. 2B), P2ry12 (Fig. 2C), Trem2 (Fig. 2D), Mafb (Fig. 2E), Cx3cr1 (Fig. 2F) and Csf1r (Fig 2G). To validate this observation, we performed immunohistochemistry (IHC) staining of P2ry12 and Tmem119 on brain slices from C57/BL6J mice that were treated with PLX diet for 2 weeks (D0) and those from Ctrl brains (Fig 2H). Iba1 was used as a microglial marker. The fluorescence intensity of P2ry12 and Tmem119 per Iba1+ cell was significantly decreased in D0 microglia compared with Ctrl microglia (Fig 2I, J).

**Fig 2.**
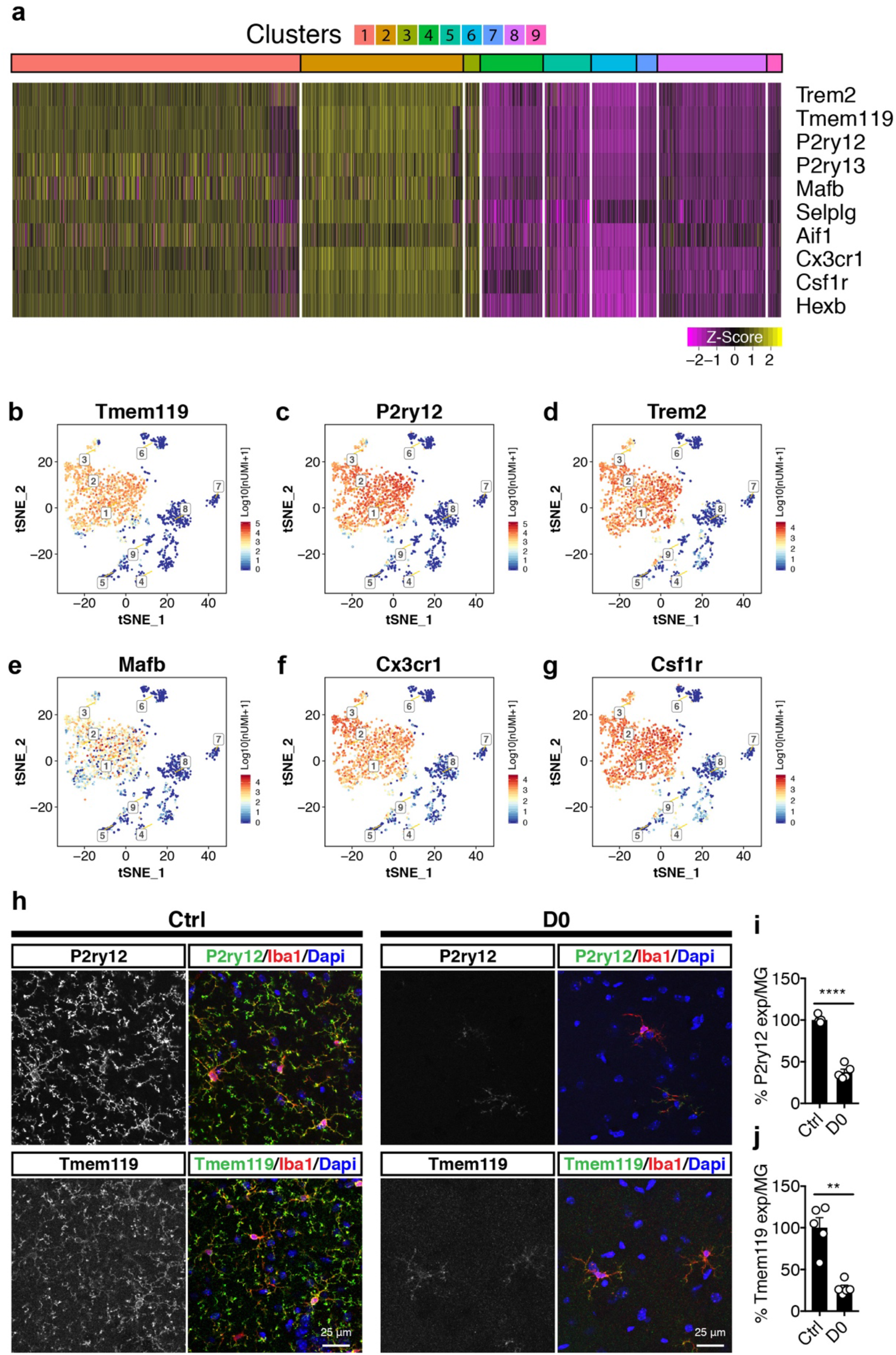
Microglial homeostatic signatures are lost in clusters that are enriched with PLX-resistant and early stage repopulating microglia. (**a**) Heatmap showing microglia homeostatic signature genes. Scaled log-transformed expression value (Z-score) was plotted. (**b-g**) tSNE plots showing expression level (log-transformed total UMI) of selected homeostatic genes: (b) Tmem119; (c) P2ry12; (d) Trem2; (e) Mafb; (f) Cx3cr1; (g) Csf1r. (**h**) Representative confocal images showing P2ry12 and Tmem119 expression in naive mice (Ctrl) and PLX treated mice (D0, 2 weeks of PLX diet). Iba1 was used as marker for microglia. Images were taken from the hippocampal region. (**i**) Quantification of relative P2ry12 expression level per microglial cell. Number of C56/BL6J mice (3-5 Mo) used: Ctrl (n=5), D0 (n=5). Unpaired t test with Welch’s correction was used. (**j**) Quantification of relative Tmem119 expression level per microglial cell. Number of C56/BL6J mice (3-5 Mo) used: Ctrl (n=5), D0 (n=5). Unpaired t test was used. P-value summary is shown as ns (p > 0.05); * (p £ 0.05); ** (p £ 0.01); *** (p ≤ 0.001); **** (p ≤ 0.0001).

### Cluster identification reveals heterogenous microglial populations under Csf1r inhibition

We next applied differentially expressed gene (DEG) analysis to obtain the transcriptomic profile of each cluster. DEGs were selected based on log fold-change ratio either above 0.5 or below −0.5 (Fig 3A and Table S1). Notably, while some of the marker genes were exclusively expressed in certain clusters, many were shared among multiple clusters. Among the most upregulated genes, each cluster could be identified with distinctive markers (Fig 3B).

**Fig 3.**
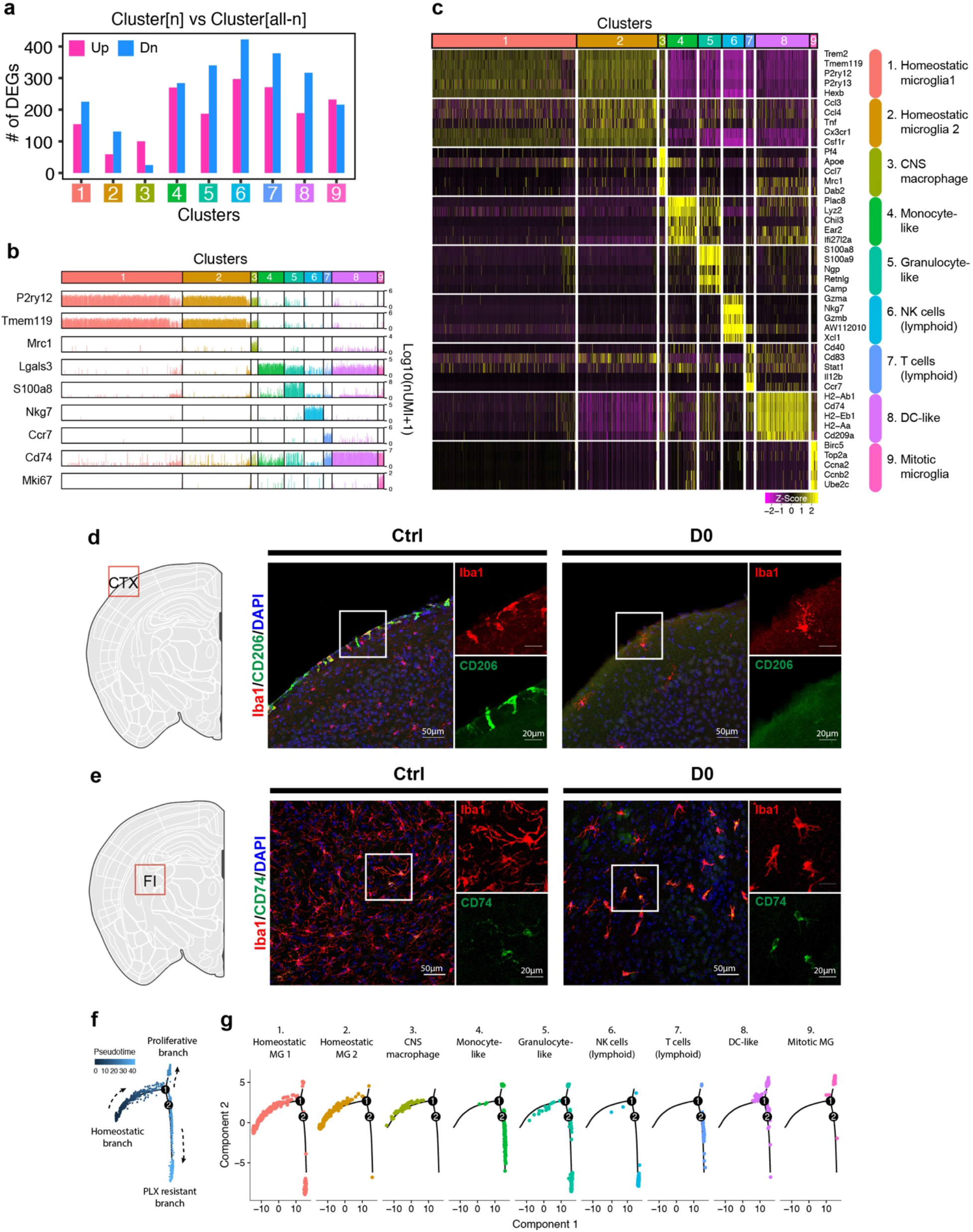
Cluster identity reveals heterogenous microglial population under Csf1r inhibition. (**a**) Bar graph showing the number of differentially expressed genes (DEGs) identified from each Cluster. Gene expression in each cluster (Cluster[n]) was compared with all the other clusters (Cluster[all-n]). (**b**) Bar graphs showing selected marker genes identified from each cluster. Log-transformed total UMI from each cell was plotted. (**c**) Heatmap showing top-5 marker genes identified from each cluster. Complete list of marker genes can be found in supplementary table 1. Annotation for each cluster is shown on the right. Scaled log-transformed expression value (Z-score) was plotted. (**d**) Representative confocal images showing immunofluorescence staining of Cd206 (green) and Iba1(red) in naïve C56/BL6J mice (Ctrl) and PLX treated mice (D0). Images were taken from the cortex (CTX). (**e**) Representative confocal images showing immunofluorescence staining of Cd74 (green) and Iba1(red) in naïve C56/BL6J mice (Ctrl) and PLX treated mice (D0). Images were taken from the fimbria (FI). (**f**) Pseudotime trajectory map that shows three different states: homeostatic, proliferative, and PLX-resistant. (**g**) Distribution of cells from each cluster placed on the pseudotime trajectory map.

In order to analyze the cell populations from each cluster in more detail, we manually compared the most prominently expressed markers from each cluster with the tabula-muris dataset, a single-cell atlas that contains annotation data for 100,000 cells from 20 different organs and tissues (Tabula Muris et al., 2018) (Fig 3C). Cluster-1 and Cluster-2 resembled homeostatic microglia, with Cluster-1 expressing high-levels of homoeostatic genes including P2ry12, Tmem119 and Trem2. Cluster-2 showed a similar homeostatic gene signature but also exhibited a modest increase of NF-kB target genes including Ccl3, Ccl4, and Tnf (Fig. 3C). Microglia in Cluster-3 expressed a high level of Mrc1, which encodes the meningeal macrophage marker CD206 (Goldmann et al., 2016), and apolipoprotein E, the major risk factor in AD. Remarkably, this population was completely wiped out by PLX treatment, as Cluster-3 was absent in both the D0 and D2 groups (Fig 1H). Immunohistochemistry analysis validated this observation, showing complete loss of CD206+ meningeal macrophages after 2 weeks of PLX treatment (Fig 3D).

Despite reduced expression of homeostatic microglial signatures, Clusters −4, −5, −8, −9 expressed myeloid markers as expected (Fig. S3A), with transcriptional profiles that resembled monocytes (Cluster-4, Fig S3B); granulocytes (Cluster-5, Fig S3B); and dendritic cells (Cluster-8, Fig S3F). Specifically, cells in Cluster-8 expressed CD209a, and MHC genes such as H2-Ab1, H2-Eb1, and CD74, a cell surface receptor for the cytokine macrophage migration inhibitory factor (MIF). Immunohistochemistry analysis showed over-representation of CD74-positive microglia in D0 brains, confirming that this cell population is resistant to csf1r inhibitors (Fig. 3E). In Cluster-9, the top-10 most upregulated genes were involved in mitosis, including Marker of Proliferation Ki-67 (Mki-67), DNA Topoisomerase 2a (Top2a), Cyclin A2 (Ccna2), and Cyclin B2 (Ccnb2) (Fig 3C). Consistent with this population representing proliferative microglia, the relative frequency of cells in Cluster 9 roughly doubled its size from 30% (D0) to 69% in just two days of microglial repopulation (Fig 1N).

Cells in Cluster-6 and Cluster-7 lacked myeloid markers including Cd11b, Csf1r, and Cd14 did not (Fig. S3A). Instead, Cluster-6 showed high expression of Klrb1c and Nkg7, which are normally found in natural killer (NK) cells (Fig S3D), while Cluster-7 expressed T-cell markers Ccr7 and Il7r (Fig S3E). We then performed microglial lineage mapping using the Cx3cr1-CreERT2 and inducible RFP reporter, and showed that microglia under Csf1r inhibition did not express either the T cell marker CD3 or the NK cell marker Nkg7 (Fig. S4). Thus, it is unlikely that these are microglia with altered states under Csf1r inhibition. Instead, it is more likely that certain subsets of NK and memory T cells were captured by cd11b-based capture during FACS since they can express Cd11b antigen as well (Chiossone et al., 2009; McFarland et al., 1992).

We then applied the pseudo-time analysis to further gain insights on the heterogeneity of microglial populations under Csf1r inhibition (Qiu et al., 2017a; Qiu et al., 2017b; Trapnell et al.,2014). By constructing an artificial trajectory tree based on the inherent gene expression data of all clusters, we revealed three branches in the pseudo-time trajectory tree: homeostatic, proliferative, and PLX-resistant (Fig 3F). As expected, Clusters −6 and −7 were placed on the PLX-resistant branch (Fig 3G), because lymphocytes were not affected by PLX5622 (Wheeler et al., 2018). Among the myeloid clusters, Cluster-8 was evenly distributed along the PLX-resistant and proliferative branches, while Clusters −4 and −5 were largely distributed along the PLX-resistant branch (Fig 3G).

### A unique Csf1r inhibition-resistant microglial population expresses Mac2

To spatially map the underlying PLX-resistant microglia from Cluster −4 and −5, we next searched for markers that distinguish them via immunohistochemistry. Marker gene analysis in Cluster-4 identified Mac2 antigen, also known as Galectin 3 (Lgals3), as a potential marker (Fig 4A). Lgals3 was highly expressed in Clusters −4, −5, −8, and −9 (Fig 4B, C). Immunofluorescence staining of brain tissues from C57/BL6J mice that were treated with PLX diet for 2 weeks (D0) revealed a subset of Iba1+ microglia expressing Mac2 in both Ctrl and D0 groups (Fig 4D), with increased Mac2 intensity and more extensive ramification in the D0 mice. After 2 weeks of PLX treatment, while the majority of Iba+ cells was depleted, showing 88% loss (Fig. 4E), the number of Iba1+Mac2+ cells was modestly increased (D0) in the hippocampus (Fig 4F). Notably, Mac2+ microglia accounted for approximately 0.68% ± 0.16 (SEM) of all Iba1+ microglia in adult hippocampus; the frequency dramatically increased to 15.46% ± 4.55 (SEM) of all remaining microglia under Csf1r inhibition (Fig 4G). The dramatic increase of the percentage of Mac2+ microglia induced by Csf1r inhibition supports the notion that this subpopulation of microglia can survive without Csf1r signaling.

**Fig 4.**
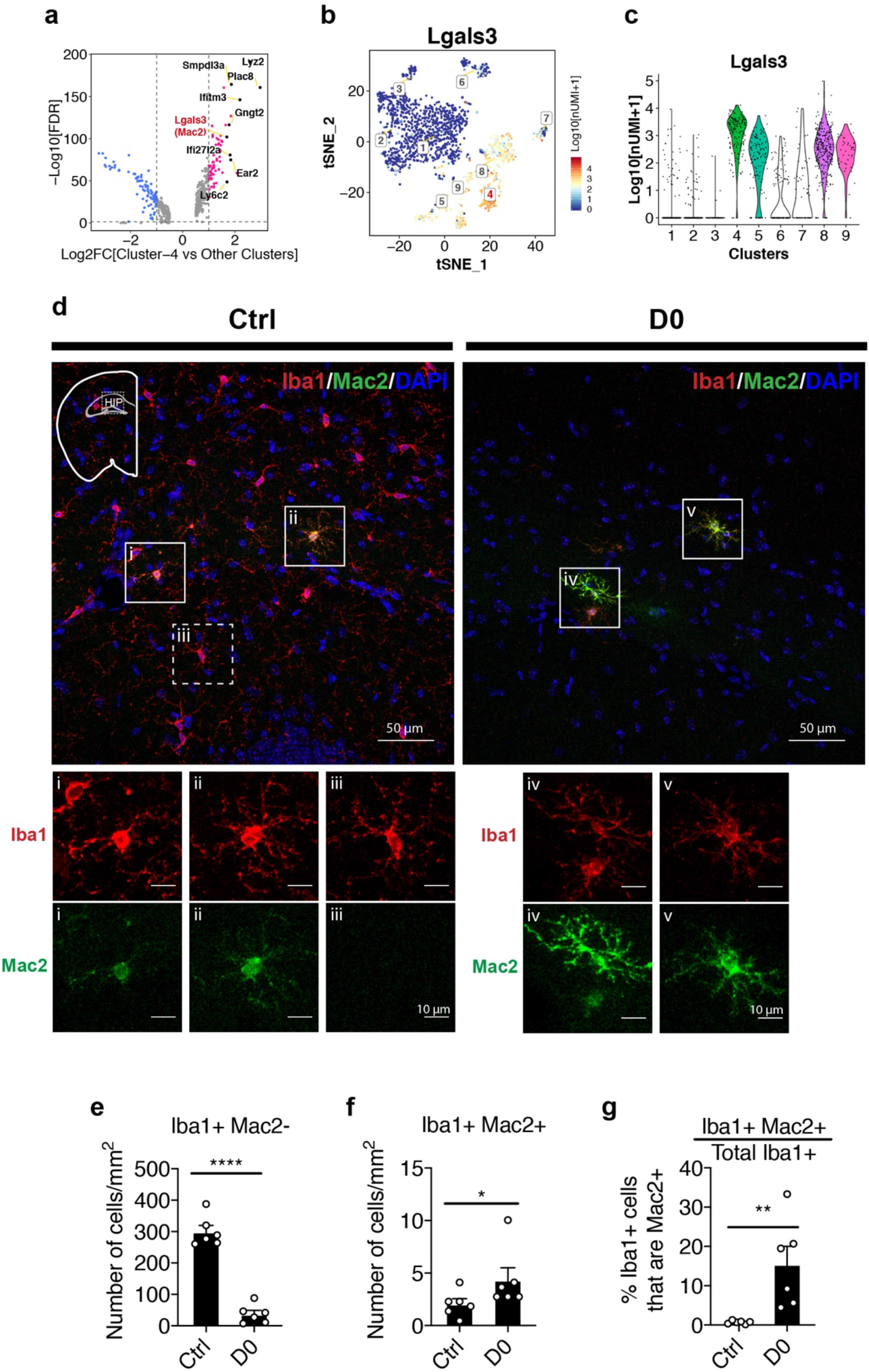
A unique Csf1 independent microglia population expresses Mac2 (Galectin-3) antigen. (**a**) Volcano plot showing differentially expressed genes identified in cluster-4. *Lgals3*, which encodes Mac2 protein is highlighted. Upregulated DEGs are colored in red and downregulated DEGs are colored in blue. (**b**) tSNE plot showing expression distribution of Lgals3 in all clusters. Log-transformed total UMI was plotted. (**c**) Violin plot showing expression of Lgals3 in all clusters. (**d**) Representative confocal images showing colocalization of Iba1 and Mac2 in native mice (Ctrl) and PLX-treated mice (D0, 2 weeks of PLX diet). Images were collected from the hippocampal region (shown in mini-map). Solid box highlights cell that is Iba1+Mac2+ and dotted box highlights cell that is Iba1+Mac2-. Enlarged images from the boxed area are shown in separate channels (panel i-v). (**e**) Quantification of Iba1+Mac2-cell numbers. Unpaired t test was used. (**f**) Quantification of Iba1+Mac2+ cell numbers. Mann Whitney test was used. (**g**) Quantification of the percentage of Mac2+ cells among all Iba1+ microglia. Unpaired t test was used. Number of C57/BL6J mice (2.5 – 4 Mo) used (panel e-g): Ctrl (n=5); D0 (n=6). Quantification in (panel e-g) was performed on images (1131.56 μm x 1938.59 μm) collected at the hippocampus using the VERSA automated slide scanner (Leica, 20x lens).

### Mac2+ microglia are not derived from peripheral monocytes and are highly proliferative

Since Mac2 is a galactoside-binding protein expressed in many myeloid cells including monocytes and macrophages (Ho and Springer, 1982), one possibility that Iba1+Mac2+ cells are over-represented under Csf1r inhibition could be due to monocytic replenishment from circulation. To test this hypothesis, we performed lineage tracing using the myeloid specific CX3CR1-CreERT2 driver (Parkhurst et al., 2013) with an inducible RFP reporter (Luche et al., 2007) (CX3CR1-CreERT2/STOP^flox^-RFP). One month after the initial tamoxifen injection, 98% labeled monocytes in circulation are replaced by unlabeled newborn cells, whereas microglia in the brain remain RFP+ due to their extreme longevity (Parkhurst et al., 2013). Using a similar labeling strategy, we examined the lineage of the parenchymal Mac2+ cells (Fig 5A). We reasoned that if the Mac2+ microglia were derived from circulating monocytes/macrophages that lack RFP expression, there would be fewer Mac2+ microglia that express RFP after repeated PLX treatment. Instead, RFP labeling efficiency in the Mac2+ population did not differ after either single (PLX^1X^) or tandem PLX treatment (PLX^2X^) separated by a repopulation period (Fig 5 B-D), indicating that the Mac2+ cells are not derived from circulating monocytes, but rather resident microglia that are internally maintained in the parenchyma.

**Fig 5.**
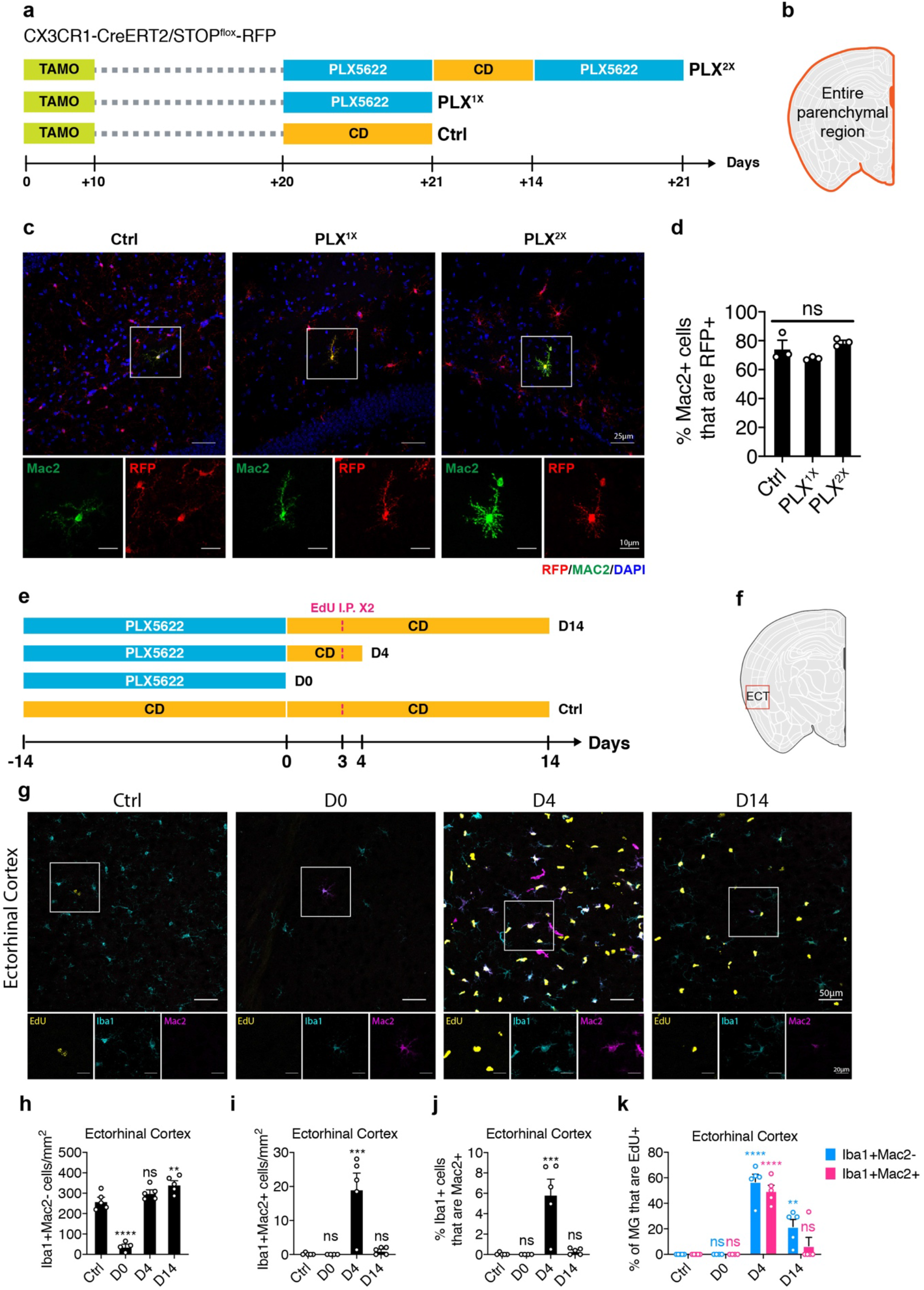
Lineage mapping shows Mac2+ microglia are not derived from circulating monocytes. (**a**) Experimental design of the lineage mapping. Cx3Cr1-CreERT2/STOP^flox^-RFP mice were injected with tamoxifen (10 days) to label microglia with RFP. Mice are either treated with PLX diet for 3 weeks (PLX^1X^) or underwent repopulation for 2 weeks and treated with PLX diet again for another 3 weeks (PLX^2X^). (**b**) Quantification area was performed on the entire parenchymal region. (**c**) Representative confocal images showing colocalization of Mac2 and RFP expression. Boxed area is enlarged and separated by each channel. Images were collected from the hippocampal region. (**d**) Quantification of the percentage of Mac2+ cell that are RFP+. Number of CX3CR1-CreERT2/STOP^flox^-RFP mice (7-9 Mo) used: Ctrl (n=3); PLX^1X^ (n=3); PLX^2X^ (n=3). One-way ANOVA was used. P-value summary is shown as ns (p > 0.05); * (p ≤ 0.05); ** (p ≤ 0.01); *** (p ≤ 0.001); **** (p ≤ 0.0001). (**e**) Experimental design of microglial repopulation timeline and EdU injections. C57/BL6J mice were treated with PLX diet for 2 weeks (D0) and switched to control diet (CD) to start repopulation for 4 days (D4) or 14 days (D14). EdU was injected on repopulation day 3. (**f**) Brain region used for quantification. Quantification in panel (h-k) was performed on images (1292.23 μm x 1130.7 μm) collected at the Ectorhinal cortex (ECT) using the VERSA automated slide scanner (Leica, 20x lens). (**g**) Representative confocal images showing immunofluorescence staining of EdU (yellow), Iba1 (cyan), and Mac2 (magenta) in the ectorhinal region. Boxed area is shown by separated channels at the bottom. (**h**) Quantification of Iba1+Mac2-cells in the ectorhinal cortex. (**i**) Quantification of Iba1+Mac2+ cells in the ectorhinal cortex. (**j**) Quantification of the percentage of Iba1+ microglia that are Mac2+ in the ectorhinal cortex. (**k**) Quantification of the percentage of EdU+ labeling in either Iba1+Mac2-cells (blue bar) or Iba1+Mac2+ cells (red bar). Number of C57/BL6 mice (2-3.5 Mo) used: Ctrl (n=5); D0 (n=4); D4 (n=5); D14 (n=5). Statistical tests used: 1) In panels (h-j), one-way ANOVA with Dunnett’s multiple comparisons test was used to compare with Ctrl; 2) In panels (k), two-way ANOVA with Dunnett’s multiple comparisons test was used to compare with Ctrl for each cell population. P-value summary is shown as ns (p > 0.05); * (p ≤ 0.05); ** (p ≤ 0.01); *** (p ≤ 0.001); **** (p ≤ 0.0001).

We then examined the newborn microglia at day 4 and day 14 after switching mice from PLX diet to the control diet using 5-Ethynyl-2’-deoxyuridine (EdU) pulse-chase labeling. To maximize the EdU labeling efficiency, two sequential intraperitoneal injections separated by 7 hours apart were given on repopulation day 3 (Fig 5E). In the ectorhinal cortex, we confirmed microglia repopulation (Fig 5F–H) and found that, while a subset of both Mac2+ and Mac2– microglia have incorporated Edu (Fig 5F, G), the number of Iba1+Mac2+ cells increased substantially at repopulation day 4 (D4) (Fig 5I), accounting for ~5% of total Iba1+ microglia (Fig 5J). Roughly 50% of the Iba1+Mac2+ cells were EdU+ at repopulation Day 4 (Fig 5K). Their highly proliferative property was further supported by the presence of mitotic marker Ki67 (Fig S5). Notably, newborn cells in the Iba1+Mac2+ population failed to retain Mac2+ expression over long term, and the number of Iba1+Mac2+ cells dropped back to control level after repopulation day 14 (Fig 5I).

### Mac2+ cells display microglial progenitor signatures

To further characterize the Mac2+ microglial population, we selected the Mac2+ cells from our scRNA-seq dataset based Lgals3 expression one standard deviation above the average log UMI counts. A total of 504 cells were manually distinguished and binned together as the Mac2+ cluster (Fig 6A). Overlay of the Mac2+ cells on the tSNE plot showed that they were largely derived from Clusters −4, −5, −8, −9, with only a few sprinkled in the homeostatic Clusters −1 and −2 (Fig 6A). To investigate the gene signatures in Mac2+ cells, we next performed DEG analysis in comparison to homeostatic Cluster-1 (Fig 3C). DEGs were identified that showed log fold-change ratio above 0.5 for upregulated (UP) genes or below −0.5 for downregulated (DN) genes. We identified a total of 385 UP and 443 DN DEGs in all Mac2+ cells (Fig 6C; Table S1). Among the DEGs, early microglial development genes such as Lyz2, Ifit3 (Matcovitch-Natan et al., 2016), Mmp8, and Mmp9 (Kierdorf et al., 2013) were significantly upregulated, while mature microglia signature genes such as Tmem119, Mafb, Cx3cr1 and Csf1r (Matcovitch-Natan et al., 2016) were downregulated (Fig 6C).

**Fig 6.**
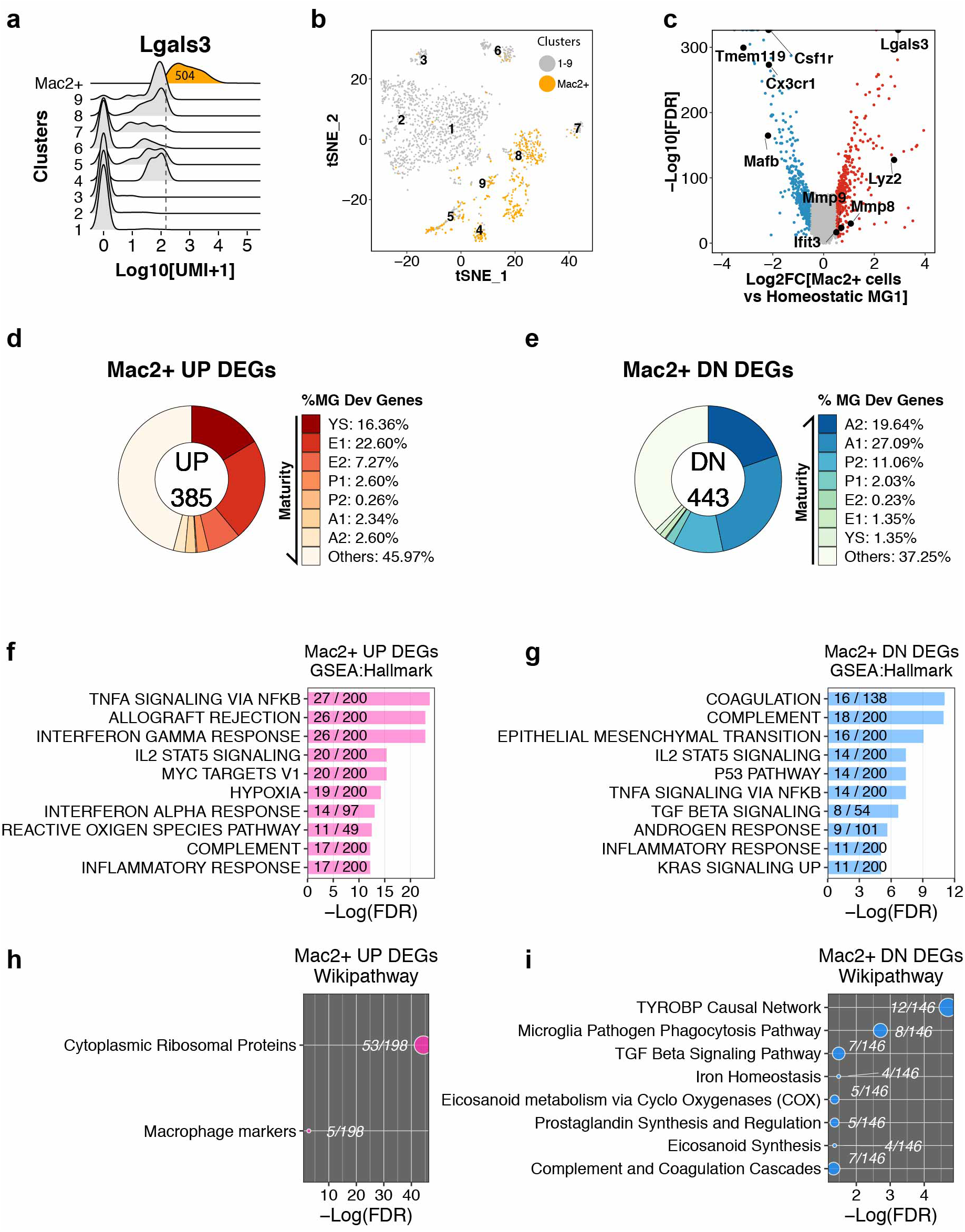
Mac2+ cells display microglial progenitor signatures. (**a**) Ridge plot showing isolation of Mac2+ cells from all clusters. Mac2+ cells were separated based on high Lgals3 expression (mean plus one SD, Log10[UMI+1] = 2.172241, shown as vertical dotted line). (**b**) tSNE plot showing the spatial distribution of Mac2+ cells in different clusters. (**c**) Volcano plot showing differentially expressed genes (DEGs) in Mac2+ cells compared to homeostatic microglia (Cluster-1). Upregulated DEGs are colored in red and downregulated DEGs are colored in blue. Genes of interest are highlighted in text. (**d**) Donut chart showing the percentage of microglial developmental genes in all upregulated DEGs in Mac2+ cells (385 genes). (**e**) Donut chart showing the percentage of microglial developmental genes in all downregulated DEGs in Mac2+ cells (443 genes). (**f**) Bar plot showing top-10 hallmark pathways enriched in upregulated Mac2+ DEGs from Gene Set Enrichment Analysis (GSEA). The fraction in the bar shows the number of genes found in the DEGs (numerator) and the number of total genes curated for the corresponding pathway (denominator). (**g**) Same analysis as in (f) but shows enrichment of pathways in downregulated DEGs. (**h**) Dot plot showing enrichment of Wikipathway terms found in upregulated DEGs in Mac2+ cells. The fraction next to the dot shows the number of genes found in the DEGs (numerator) and the number of total genes curated in the corresponding pathway (denominator). (**i**) Same analysis as in (h) but shows enrichment of pathways found in downregulated DEGs.

To further examine whether the Mac2+ cells resembled immature microglial progenitors during development, we compared the DEGs of Mac2+ cells with the microglial developmental gene sets identified previously (Matcovitch-Natan et al., 2016). In keeping with the original terminology, a total of 7 different gene clusters were compared, which covered the entire microglial developmental trajectory, from yolk sac (Cluster YS, day E10.5 – E12.5), early microglia (Cluster E1/E2, day E10.5 – E14), pre-microglia (Cluster P1/P2, day E14 to postnatal day 9), to adulthood (Cluster A1/A2, >1Mo) (Matcovitch-Natan et al., 2016). Remarkably, genes associated with early development were highly enriched among the upregulated DEGs in the Mac2+ cells (Fig 6D). Specifically, 16.36% of genes were those associated with yolk sac (YS) progenitors and 22.60% of the genes were those expressed in early embryonic progenitors (E1), while only around 2% of those expressed in adult microglia (Fig 6D). In contrast, among the downregulated DEGs from Mac2+ cells, 19.64% and 27.09% of them overlapped with the A2 and A1 adult gene signatures respectively (Fig. 6E). As a control for the analysis, we also examined the cells lacking Mac2 expression (Mac2–), which exhibit no progenitor signatures in them (Fig S6). In direct contrast to Mac2+ cells, upregulated DEGs in Mac2-cells showed significant overlap with adult microglia while the downregulated one with yolk sac (Fig S6).

We next performed Gene Set Enrichment Analysis (GSEA) of the DEGs in the Mac2+ microglia (Mootha et al., 2003; Subramanian et al., 2005). NF-kb pathway and interferon signaling were highly enriched in the upregulated DEGs (Fig 6F), whereas p53, STAT5, and TGFbeta signaling, a pathway required for microglia homeostasis maintenance (Butovsky et al., 2014; Zoller et al., 2018), was enriched among the downregulated DEGs (Fig 6G). Additional analyses with Wikipathway (Slenter et al., 2018) showed ribosomal protein and macrophage signature genes in the upregulated DEGs, whereas the Tyrobp network and pathogen phagocytosis pathway were the most enriched signature among the downregulated DEGs. As Tyrobp is the downstream adaptor of Csf1r signaling (Otero et al., 2009), its downregulation could explain the resistance of Mac2+ cells to Csf1r inhibition.

### Mac2+ progenitor-like cells are present among homeostatic microglia

We next questioned whether the Mac2+ cells in non-treated ctrl brain also have progenitor-like features. Among all the Mac2+ cell from our scRNA-seq data, 3.01% came from the untreated ctrl C57/BL6J mice (Fig 7A). Mac2+ cells from Ctrl mice exhibited similar upregulation of developmental genes but downregulation of homeostatic markers compared to cluster-1 (Fig 7B). Specifically, approximately 86.44% upregulated DEGs (Fig 7C) and 93.67% downregulated DEGs (Fig 7D) from the Mac2+ cells in Ctrl brains were found in all Mac2+ cells, suggesting that the properties of Mac2+ cells in the presence or absence of csf1r inhibition are indistinguishable. Indeed, Mac2+ cells from the control brain exhibited similar microglial progenitor signatures with reduced expression of mature microglial markers (Fig 7E, F), further confirming the presence of Mac2+ progenitor-like microglia in adult mouse brain.

**Fig 7.**
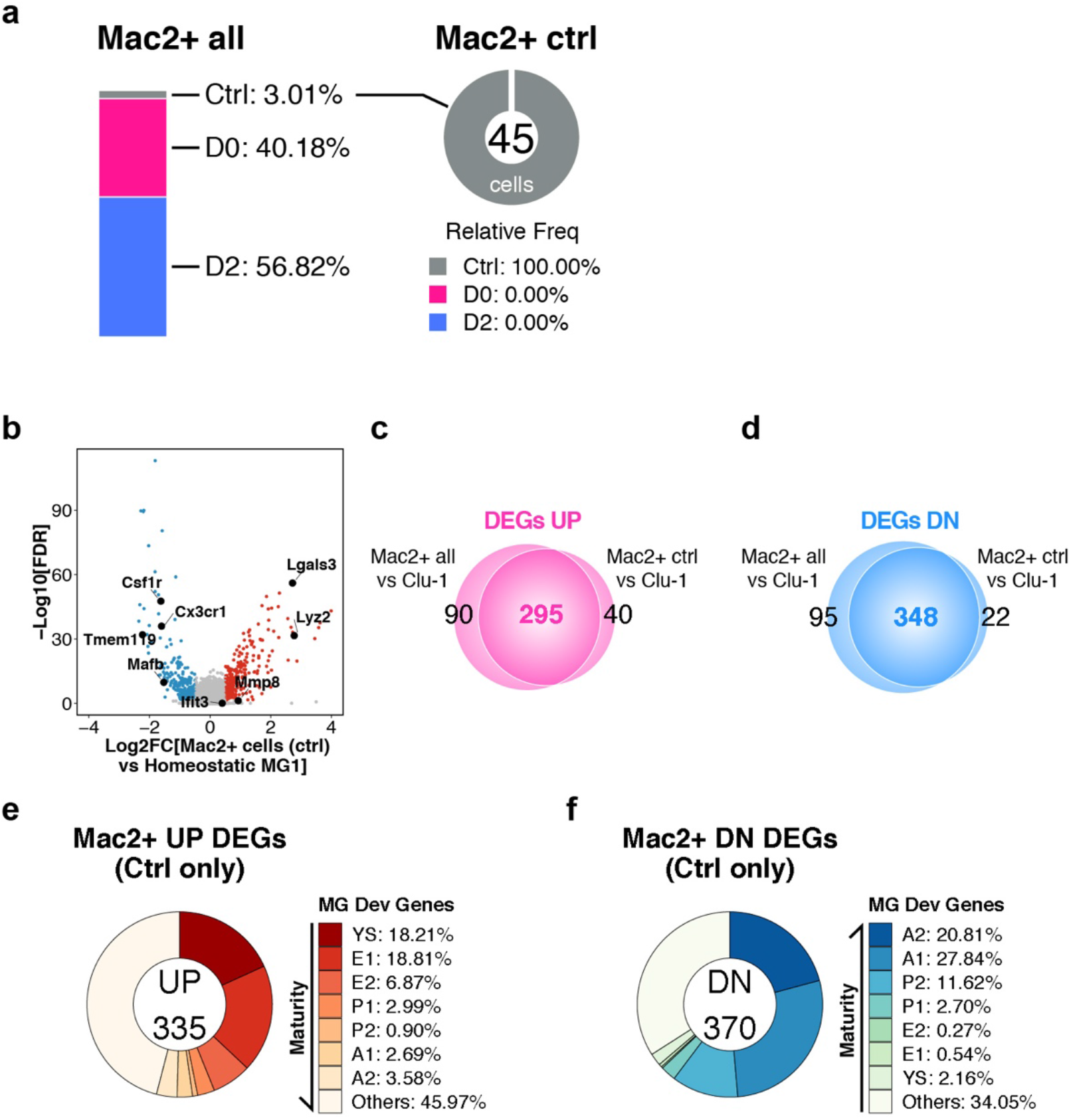
Mac2+ progenitor-like cells are present among homeostatic microglia. (**a**) Bar graph showing the relative frequency of cells from each treatment group distributed in all Mac2+ cells. A total 45 Mac2+cells were from homeostatic ctrl brain (3.01%). (**b**) Volcano plot showing differentially expressed genes (DEGs) in Mac2+ ctrl cells in comparison to homeostatic microglia (cluster-1). Upregulated DEGs are colored in red while downregulated DEGs are colored in blue. Genes of interest are highlighted in text. (**c**) Venn diagram showing the common upregulated DEGs found between all Mac2+ cells (left circle) and Mac2+ cells from ctrl brain (right circle). (**d**) Venn diagram showing the common downregulated DEGs found between all Mac2+ cells (left circle) and Mac2+ cells from ctrl brain (right circle). (**e**) Donut chart showing the percentage of microglial developmental genes among the upregulated DEGs in Mac2+ cells from ctrl brain (335 genes). (**f**) Donut chart showing the percentage of microglial developmental genes among the downregulated DEGs in Mac2+ cells from ctrl brain (370 genes).

## DISCUSSION

In the current study, we employed scRNA-seq to characterize the CNS myeloid population under acute Csf1r inhibition and identified a resistant microglia population that expresses Mac2 antigen. In the hippocampus, the Mac2+ microglial sub-population represented 0.68% total microglia and did not require Csf1r signaling for survival. While the Mac2+ microglia shared transcriptomic features similar to those of circulating monocytes, lineage tracing revealed that the Mac2+ microglial population was self-sustained with no replenishment from circulation. Remarkably, the Mac2+ population shared striking similarities with immature microglial progenitors during development. Finally, Mac2+ microglia appeared to be highly proliferative in the ectorhinal region during adult microglial repopulation after acute ablation. Altogether, our data identify a heterogeneous progenitor-like microglial population that is independent of Csf1r survival signal in adult mouse brain.

The existence of a Csf1r-independent microglial population in the adult CNS was suggested by previous studies. For example, regardless of the dose and duration of PLX treatment, 1-10% of the microglial population has been observed to survive (Acharya et al., 2016; Huang et al., 2018; Rice et al., 2017; Zhan et al., 2019). In Csf1r KO mice, although 99% of Iba1+ microglia are lost in almost all brain regions, 13.9% microglia remain in the hippocampus and 33.7% remain in the piriform cortex (Erblich et al., 2011). During development, although it is well known that myeloid cells require Csf1r signaling for survival, the CD45+c-kit^lo^ microglial progenitors which are found in yolk sac do not start to massively express Csf1r until days post coitum (dpc) 9.0 (Kierdorf et al., 2013). Furthermore, expression of Csf1 ligand remains mostly quiescent until yolk sac progenitors reach the brain rudiment at E10.5 (Matcovitch-Natan et al., 2016). These findings support the notion that a Csf1r-independent microglial population in the adult CNS might reflect a similar paradigm to the developmental progenitors in which alternative survival pathways are engaged.

In characterizing the remaining microglia after acute Csf1r inhibition, we recovered a subset of cells that can be distinguished by the Mac2 antigen that was also detectable under homeostatic conditions. Unlike the vast majority of microglial population that is sensitive to Csf1r inhibitor, the Mac2+ population modestly increased in response to the drug (Fig 4F). The Mac2+ cells uncovered under either homeostasis (Ctrl) or under Csf1r inhibition/early repopulation (D0/D2) exhibited striking similarities with early microglial progenitors. In particular, Mmp8 and Mmp9, which are required for the migratory expansion during early stage microgliogenesis (Kierdorf et al., 2013) were highly upregulated whereas Mafb, a transcriptional factor required for microglia maturation (Matcovitch-Natan et al., 2016) was significantly downregulated. Comprehensive comparison of transcriptomes revealed that Mac2+ microglia in adult brain exhibit significant overlap with those found in yolk sac and in early embryonic development, highlighting the unusual heterogeneity and plasticity of adult microglia.

Mac2, also known as Galectin-3, is a galactoside binding protein highly expressed in myeloid cells that can be secreted to modulate a wide variety of immune functions (Rahimian et al., 2018). Using fate-mapping, we determined that the Mac2+ cells observed under Csf1r inhibition were not of monocytic origin, but represent a subpopulation of microglia. Expression of Mac2 is associated with activated microglial state (Lalancette-Hebert et al., 2012). Mac2 is found to promote microglial migration (Wesley et al., 2013). Microglia in Mac2 knockout mice have far less proliferation in response to ischemic lesions (Lalancette-Hebert et al., 2012), suggesting a critical role of Mac2 in modulation of microglial proliferation. Whether depletion of Mac2 diminishes the repopulation microglia following Csf1r inhibition remains to be determined. Interestingly, a more recent study showed Mac2 colocalizes with Trem2 in microglial processes, and stimulates Trem2-Tyrobp signaling (Boza-Serrano et al., 2019). Whether activation of Trem2-Tyrobp signaling is involved in the resistance of Mac2+ microglia to Csf1r inhibitor is not known.

Our findings revealed a progenitor population hidden among other microglia under steady-state. Unlike other terminally differentiated myeloid cells, microglia are sufficiently self-maintained (Ajami et al., 2007; Mildner et al., 2007). Under acute microglial ablation, the 1-10% remaining microglia can restore the entire empty niche (Huang et al., 2018; Zhan et al., 2019), without needing any external progenitor input. However, due to the unusual plasticity of microglia, it still remains unclear whether a specialized adult progenitor is required to maintain microglial homeostasis. It had been proposed that every microglial cell could have the ability to conduct self-renewal when needed (Tay et al., 2017). Consistent with this notion, although we found that the Mac2+ cells were highly proliferative during early stage microglial repopulation, a large proportion of proliferating microglia did not express Mac2. And the number of Mac2+ cells returned to ctrl level by D14. The overall contribution of Mac2+ microglia during repopulation is hard to determine with the current approach since Mac2 expression could be very transient, and rapidly turned off once the cell cycle is complete. To determine the overall contribution of Mac2+ cells to microgliogenesis, further lineage tracing studies of the descendants of Mac2+ progenitors are needed.

## METHODS

### Mice

All animal work was performed in accordance with the Institutional Animal Care and Use Committee guidelines, at the University of California, San Francisco. Mice, with unrestricted access to water and food source, were housed in a pathogen-free barrier facility operated on a 12 hr light on/off cycle. The C57BL/6J mice were supplied by National Institute on Aging (Charles River, Wilmington, MA, USA). CX3CR1-CreERT2/STOP^Flox^-RFP mice were generated by crossing the CX3CR1-CreERT2 line (JAX: 021160) and the STOP^Flox^-RFP line (MGI: 104735). Equal numbers of male and female mice were used for all experiments except for the sc-RNAseq experiments, which used only female mice.

### Drug administrations

For acute microglial ablation, mice were administered Csf1r antagonist PLX5622 orally via PLX diet (1200 mg/kg PLX5622, Plexxikon Inc., Berkeley, USA). Control diet (CD) with the same base formula was used as control. For lineage mapping, tamoxifen (Sigma-Aldrich, T5648) was dissolved in corn oil and administered to the CX3CR1-CreERT2/STOP^Flox^-RFP mice by intraperitoneal (I.P.) injection for 10 days at a daily dose of 2 mg. To label proliferative cells, a solution containing 20 mg/mL 5-Ethynyl-2’-deoxyuridine (EdU) (Santa Cruz, sc-284628) was prepared fresh in sterile PBS and given to mice via I.P. injection at a dose of 80 mg/kg per animal. To maximize EdU labeling, two injections, separated by 7 hours, were given on the same day (Repopulation day 3). To detect EdU-labeled cells in brain sections, Click-iT™ EdU imaging kits (ThermoFisher Scientific, C10337) were used following the manufacturer’s instructions before the immunofluorescence staining procedures.

### Tissue preparation for microglia isolation

Four female mice (1 for Ctrl, 2 for D0, 1 for D2) were perfused with PBS transcardially to remove circulating blood cells in the CNS. Whole brain was then dissected, with the cerebellum removed, and homogenized in an enzymatic digestion buffer containing 0.2% Collagenase Type 3 (Worthington, LS004182) and 3 U/mL Dispase (Worthington, LS02104). The digestion was performed at 37 °C for 45 min and quenched by an inactivation buffer containing 2.5 mM EDTA (Thermofisher, 15575020) and 1% fetal bovine serum (Invitrogen, 10082147). The homogenates were kept at 4°C for all downstream applications. The homogenates were then processed for myelin depletion using myelin removal beads (Miltenyi Biotec, 130-096-733) and passed through LD column (Miltenyi Biotec, 130-042-901). The myelin-depleted fraction was then used for FACS.

### Adult microglia purification via fluorescence-activated cell sorting (FACS)

To perform immune labeling, myelin-depleted cell suspensions were incubated with TruStain fcX (BioLegend, Cat. No. 101319, clone 93) for 10 min at 4°C to block Fc receptors (1:50 dilution). To label myeloid cells, the homogenates were incubated with a Cd11b antibody conjugated with APC fluorophore for 20 min at 4°C (Tonbo bioscience, Cat: 20-0112, clone M1/70, 1:100 dilution). Sytox-blue live/dead stain (Thermofisher, S34857) was included 5 min before sorting (1:1000 dilution). Cell suspensions were then sorted on a flow cytometer (BD FACSAria II). Live Cd11b+ fraction was then sorted into pre-chilled RPMI-1640 media (Thermofisher Scientific, 12633012) containing 5% FBS. FACS gating strategy is shown in Supplementary Fig 1. The FACS procedures were performed by the Gladstone Flow Core (Gladstone Institutes, San Francisco, USA).

### Single-cell cDNA library preparation and sequencing

Single-cell gene expression profiling was performed using the 10x Genomics platform according to the manufacturer’s instructions (10x Genomics, Pleasanton, CA, USA). FACS sorted cells were captured in droplets that were emulsified with gel beads containing barcoded primers (https://www.nature.com/articles/ncomms14049). Single-cell expression libraries were prepared using the 10x genomics Chromium single cell 3′ library and gel bead kit v2 reagents (10x Genomics, PN-120237). To minimize batch effects, all 4 samples were prepared simultaneously in a single assay. Briefly, suspension containing 2,000-10,000 single cells were loaded into the 10x single cell A chip and processed in the 10x Chromium controller. After reverse transcription, cDNAs were amplified by 12-14 PCR cycles. An aliquot of 35 ul cDNA product was then used for library construction, which included 14 cycles of sample index PCR to barcode samples. The quality of the cDNA library was checked by the Bioanalyzer (Agilent). KAPA qPCR was performed to measure the library quantities. A pool of all quantity-balanced libraries was then sequenced on the NextSeq 500 sequencer (Illumina) at an average sequencing depth of 45,002 reads/cell (post normalization). The single-cell library preparation and sequencing steps were performed by the Gladstone Genomics Core (Gladstone Institutes, San Francisco, USA).

### Bioinformatics

The Illumina BCL output files were processed by the Cell Ranger software (2.0.1, 10x Genomics). Sequencing reads were mapped to the mouse genome (mm10). A total of 2,149 cells were captured. On average, 1,607 genes/cell and 4,632 UMIs/cell were detected. Over 94.6% reads were detected in cells. Downstream gene profile analyses were performed using the Seurat package in R [Seurat 2.0, (Butler et al., 2018)]. The data set was filtered based on the following criteria: 1) cells with UMI count below 500 or above 20,000 were removed; 2) cells that have less than 200 genes were removed; 3) cells that have greater than 10% mitochondrial genes were removed; (Supplementary figure 2 a-d). In addition, we also removed genes that showed expression in less than 10 cells. After data filtering, we obtained a total 2,100 cells for subsequent analysis (Fig 1c). The data were normalized by log transformation followed by regression based on total UMI counts and mitochondrial gene content. Genes associated with principle components (1 to 22) were used for data dimensional reduction based on tSNE to generate distinctive cell clusters (Supplementary figure 2e, f). Data visualization including heatmaps, tSNE plots, and violin plots, ridge plots were generated using the following built-in functions from Seurat 2.0: “DoHeatmap”, “FeaturePlot”, “VlnPlot”, and “RidgePlot”. Differentially expressed genes (DEGs) were performed using the “FindMarkers” function with log fold-change threshold set at 0.5. GLM-framework Mast was used as statistical test for the DEG analysis (Finak et al., 2015). Cluster annotation was performed manually with reference to the Tabula-muris dataset (Tabula Muris et al., 2018). Pseudo-time analysis was performed using the monocle package in R [Monocle 2.0, (Qiu et al., 2017a; Qiu et al., 2017b; Trapnell et al., 2014)]. Gene network analyses were performed with gene set enrichment analysis (GESA) with molecular signatures database (MSigDB) (Mootha et al., 2003; Subramanian et al., 2005). Pathway analysis was performed using the Bioconductor R package “rWikiPathways” (Slenter et al., 2018). The raw count matrix of the scRNA-seq data can be access on GEO with accession number (GSE133773). A list of R codes used for the analyses are available on Github (https://github.com/lihong1github/Zhan-et-al-2019-scRNAseq).

### Immunohistochemistry

Perfused mouse brains were fixed in 4% paraformaldehyde prepared in PBS for 48 hr and then incubated in 30% sucrose for at least another 48 hr. Coronal sections of 30 μm thickness were obtained by cutting the fixed brains on a sliding microtome (Leica, SM2010R). Sections were stored in cyroprotectant and kept at −20 °C before use. To perform immunohistochemistry staining, sections near the same stereological position were used. One or two sections per mouse were used for each staining. Floating sections were washed in PBST buffer (PBS containing 0.5% Triton X-100) 3 times for 5 mins each. 3% normal donkey serum (NDS) was used for blocking at room-temperature for 1 hr. Floating sections were incubated with primary antibodies diluted in PBST containing 3% NDS at 4 °C overnight. The following primary antibodies and respective dilution ratio were used: P2ry12 (1:500, a gift from Dr. David Julius, University of California San Francisco); Tmem119 (1:250, Abcam, ab209064); Iba1 (1:500, Abcam, ab5076); Mac2 (1:1000, Cedarlane, CL8942AP); RFP (1:500 Rockland, 200-101-379); S100a8 (Thermo fisher, PA5-79948); Cd206 (1:500, Abcam, ab64693); Cd74 (1:500, Abcam, ab245692); Cd3 (1:250, Abcam, ab16669); Nkg7 (1:50, Thermo fisher, PA5-67173); Ki67 (1:100, Abcam, ab16667). Antigen retrieval was performed for Ki67 staining. For antigen retrieval, floating sections were incubated in sodium citrate buffer (10 mM sodium citrate, 0.05% Tween 20, pH 6.0) at 95°C for 30 min before blocking.

Secondary antibodies were diluted at 1:1000 in PBST containing 3% NDS. Floating sections were incubated with diluted secondary antibodies at room temperature for 1hr with gentle shaking. All secondary antibodies used were donkey IgG conjugated with different fluorophores including, Cy3, Alexa Fluor 488, Alexa Fluor 647. All secondary antibodies were obtained from Jackson ImmunoResearch. After secondary antibody staining, sections were washed in PBST for 4 times, 10 min each. DAPI was included in the last washing step as nuclei stain. Sections were then placed on glass slides and mounted with VECTASHIELD antifade mounting media (Vector Laboratories, H-1000).

### Microscopy

Immunofluorescence images of the sections were acquired on the VERSA automated slide scanner (Leica Biosystems, Wetzlar, Germany). The microscope was equipped with an Andor Zyla 5.5 sCMOS camera (Andor Technologies, Belfast, UK) and was funded under NIH S10 grant OD021717. Image acquisition was performed using the ImageScope software (Aperio Technologies, Vista, CA). The output SCN image files were processed by the Bio-Formats software (Linkert et al., 2010). Confocal microscopy was performed on a Zeiss LSM880 inverted scanning confocal microscope (Carl Zeiss Microscopy, Thornwood, NY). The microscope is equipped with 2 PMT detectors, a high-sensitivity GaAsP detector, and a 32-GaAsP Airyscan super resolution detector. Images were acquired using the Zeiss Zen imaging software. Z-stacks of confocal images were acquired with 8-12 focal planes at 0.75-1 μm interval. Representative images were created using max projections from Z-stacks.

### Image analyses and quantification

All image analyses were performed in FIJI V1.50i (Schindelin et al., 2012). Analysis macros were written using the IJM scripting. Briefly, multi-channel images were split to each induvial channel. Image segmentation was performed using the adaptive threshold approach (https://sites.google.com/site/qingzongtseng/adaptivethreshold). Cells showing two different markers (e.g. Iba1+ and Mac2+) were selected if the cell mask from each channel showed greater than 10% overlap. Quantification including cell counting and area measurement was performed using the “Analyze Particles” and “ROI Manager” function in FIJI. A list of the analysis IJM code used in the study are available on Github (https://github.com/lihong1github/Zhan-et-al-2019-scRNAseq).

### Statistics

All experiments were performed with a minimum of at least 3 biological replicates. Mean values from each animal were used for computing statistical differences. Standard error of the mean (SEM) was used for error bars. Statistical analyses were performed in Graphpad prism 8.0 (Graphpad, San Diego, CA) and R (F Foundation for Statistical Computing, Vienna, Austria). Data visualization was achieved with R package ggplot2 (Wickham, 2009). Data normality was assessed using the Shapiro-Wilk normality test. F test was used to assess homoscedasticity prior to unpaired t-test and Brown-Forsythe test was used to homoscedasticity prior to ANOVA. For data with normal distribution and equal variance, unpaired t-test was used to compare two groups. One-way ANOVA was used to compare data with more than two groups. Dunnett’s multiple comparisons test was used to compare difference between designated groups. Two-way ANOVA with Dunnett’s multiple comparisons test was used for multiple group comparison to Ctrl. For data that failed to pass the normality test, Mann Whitney test was applied. For data with unequal variance, unpaired t-test with Welch’s correction was applied. P-value and FDR are summarized as ns (p > 0.05); * (p ≤ 0.05); ** (p ≤ 0.01); *** (p ≤ 0.001); **** (p £ 0.0001).

## ACKNOWLEDGEMENTS

We want to thank Dr. David Julius from University of California San Francisco for the P2ry12 antibody; Parmveer Singh from Plexxikon Inc., for supplying the PLX diet; Nandhini Raman from Gladstone Flow Cytometry Core for FACS assistance; Jim McGuire and Dr. Natasha Carli from the Gladstone Genomics Core for performing single cell RNA library preparation and sequencing QCs procedures; Steve Belunek from Gladstone IT for assistance on cluster computing; Dr. Matthew Lee Settles from University of California-Davis for the scRNA-seq analysis training course; Cody Jackson from the Gladstone Histology and Light Microscopy Core for imaging assistance; and Dr. Kathryn Claiborn for editing the manuscript. NIH S10 RR028962 funded the use of FACSAria cell sorter; James B. Pendleton Charitable Trust funded the use of NextSeq 500 sequencer. This work was supported by NIH grants 1R01AG054214-01A1, U54NS100717, R01AG051390, and Tau Consortium grant (to L.G.)

## AUTHOR CONTRIBUTIONS

L.Z. and L.G. conceived the project and designed the experiments. L.Z., P.D.S., Y.Z., and Y.L. performed the experiments and analyzed the data. L.Z., P.D.S., and L.G. wrote the manuscript.

