## Supplemental Figures for "A Mac2-positive progenitor-like microglial population survives independent of CSF1R signaling in adult mouse brain"

**Supplemental information**

Supplementary data include 6 figures and 1 table.

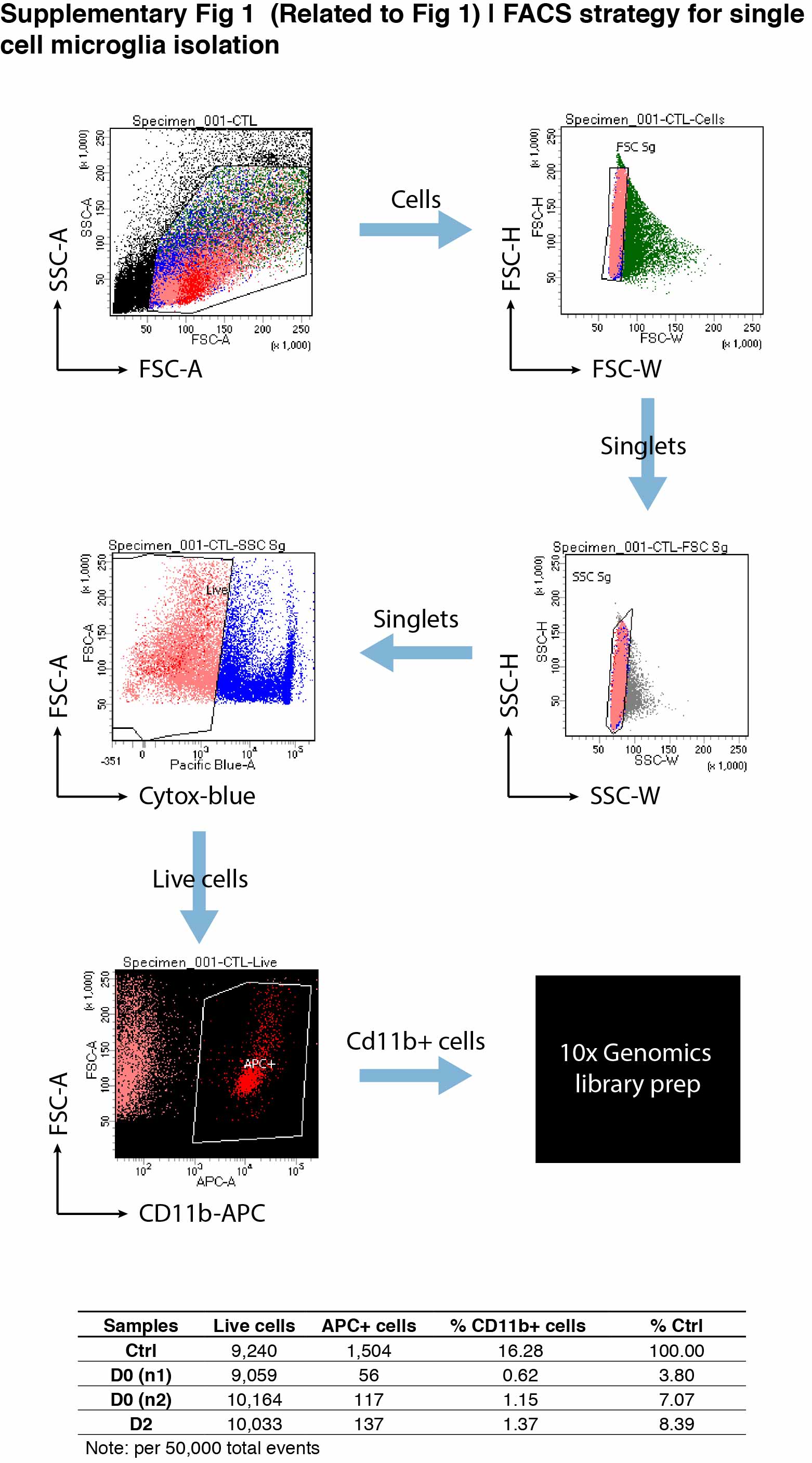

**Supplementary Fig 1 (related to Fig 1) | FACS strategy for single cell microglia isolation.**

Gating plots used for cell sorting are shown. Plots were generated using BD FACSDiva software. The table at bottom shows the summary from 50,000 events recorded.

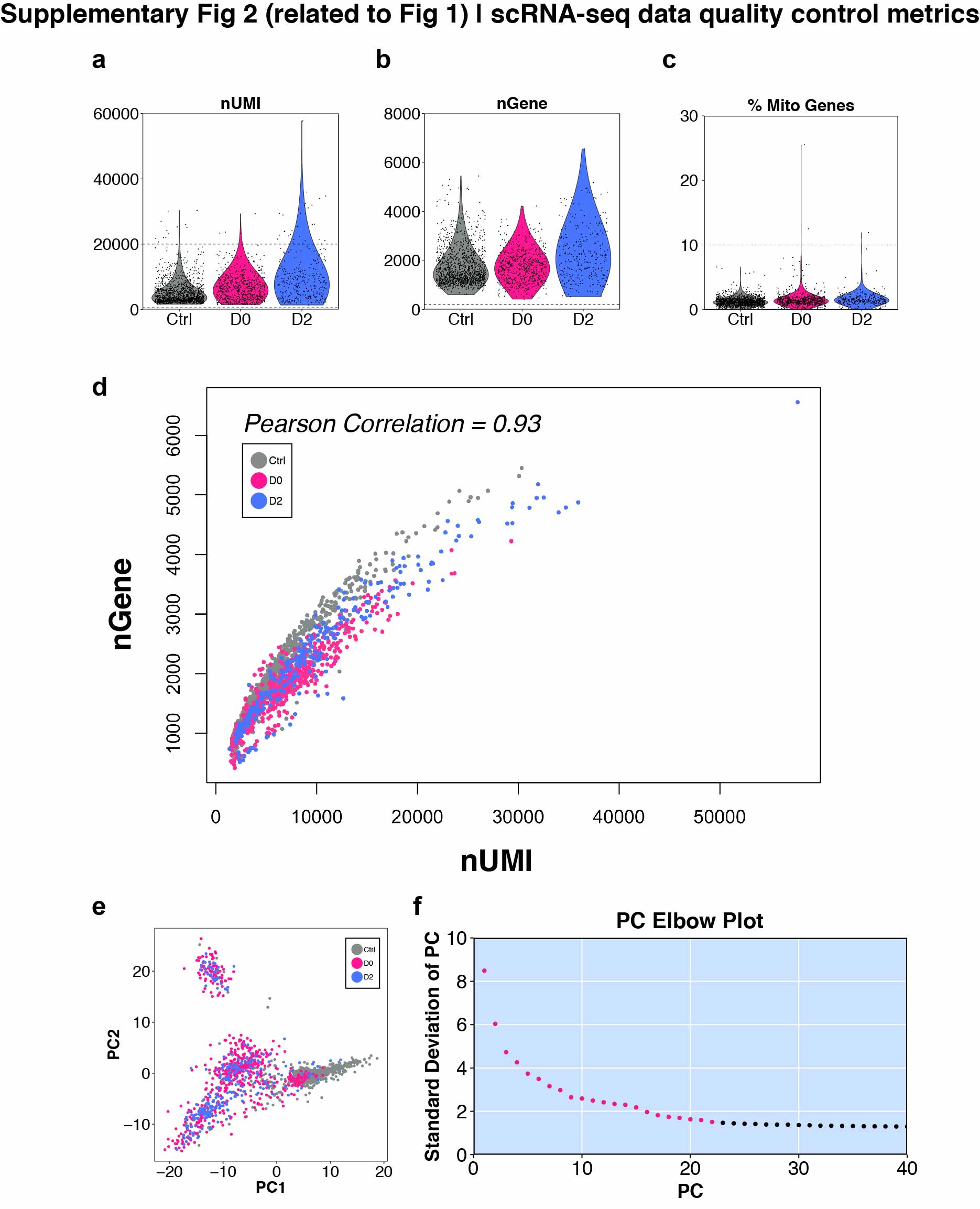

**Supplementary Fig 2 (related to Fig 1) | scRNA-seq data quality control metrics**

(**a**) Violin plot showing UMI counts for each treatment group. Dotted line shows the cell filter cut-off. (**b**) Violin plot showing total number of genes for each treatment group. Dotted line shows the cell filter cut-off. (**c**) Violin plot showing the parentage of mitochondrial genes per cell for each treatment group. Dotted line shows the cell filter cut-off. (**d**) Scatter plot showing the correlation between total number of genes detected and total number of UMI. Pearson correlation (0.93) is shown. (**e**) Scatter plot from principle component analysis (PCA). PC1 and PC2 are shown. (**f**) Elbow plot showing standard deviation of the first 40 principle components from PCA. Principle components 1 to 22 (red colored) were used as input for downstream tSNE analysis.

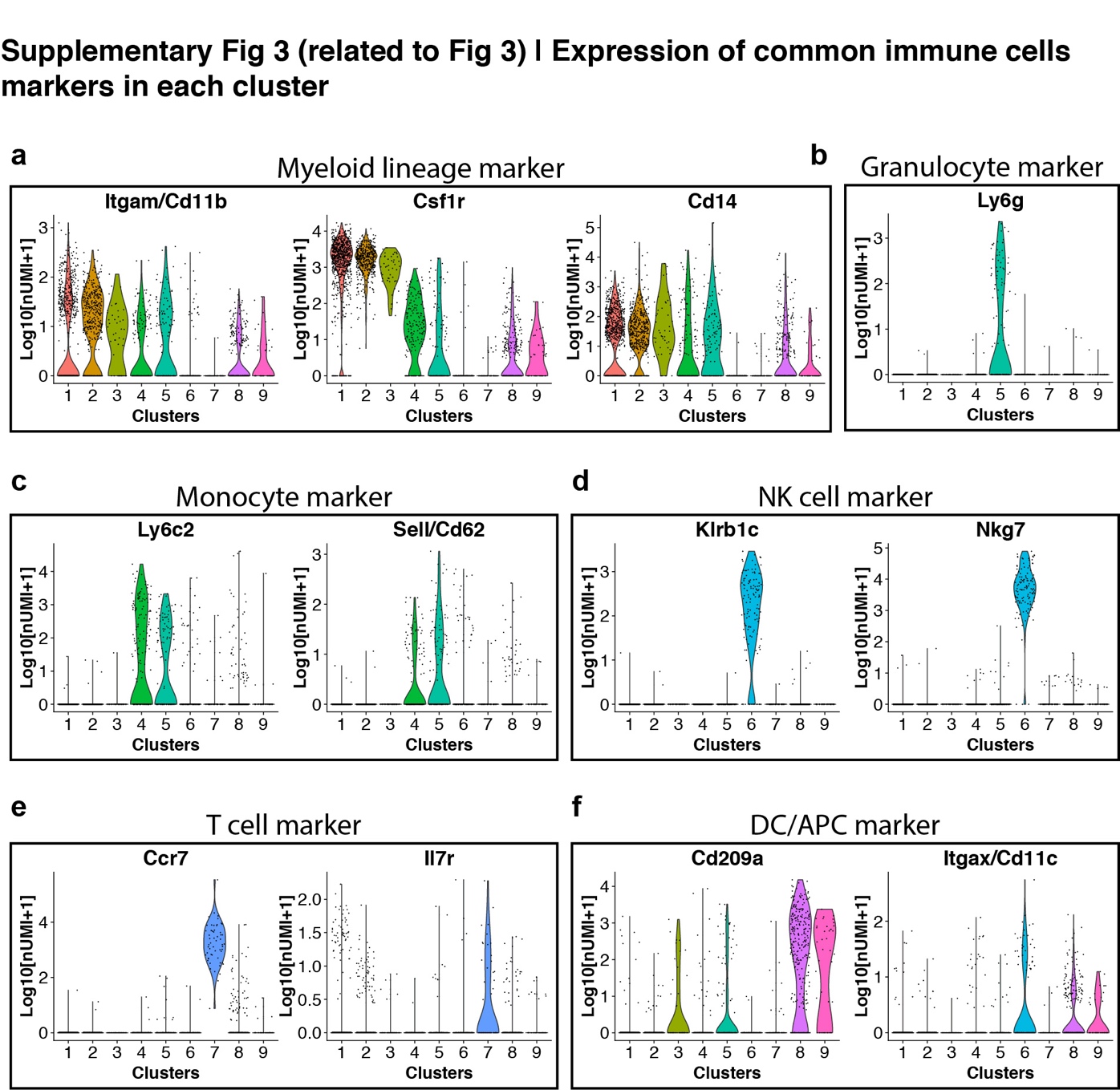

**Supplementary Fig 3 (related to Fig 3) | Expression of common immune cells markers in each cluster**

(**a**) Violin plots showing expression of myeloid marker Itgam, Csf1r, Cd14 in all clusters. (**b**) Violin plot showing expression of granulocyte marker Ly6g. (**c**) Violin plots showing expression of monocyte marker Ly6c2 and Sell (Cd62). (**d**) Violin plots showing expression of NK cell marker Klrb1c and Nkg7. (**e**) Violin plots showing expression of T cell marker Ccr7 and Il7r. (**f**) Violin plots showing expression of dendritic cell (DC) marker Cd209a and Itgax (Cd11c).

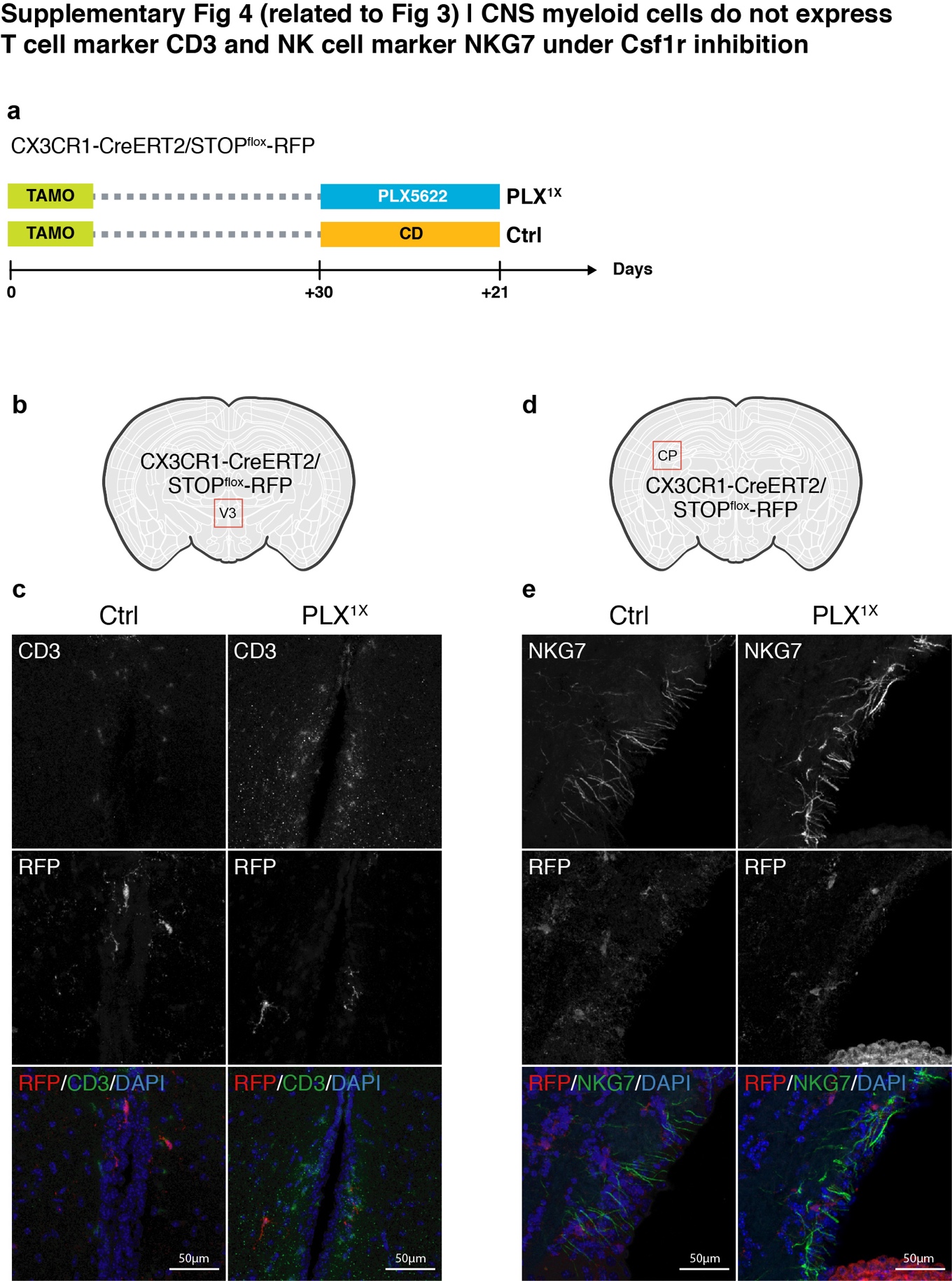

**Supplementary Fig 4 (related to Fig 3) | CNS myeloid cells do not express T cell marker CD3 and NK cell marker NKG7 under Csf1r inhibition** (**a**) Experimental design of the lineage mapping. Cx3Cr1-CreERT2/STOP^flox^-RFP mice were injected with tamoxifen (10 days) to label microglia with RFP. Mice are either treated with PLX diet for 3 weeks (PLX^1X^). (**b**) Imaging region for T cell marker Cd3 at 3^rd^ ventricle (V3) in CX3CR1-CreERT2/STOP^flox^-RFP mice. (**c**) Representative confocal images showing immunofluorescence staining of Cd3 and RFP in untreated control mice and PLX treated mice (PLX^1X^). (**d**) Imaging region for NK cell marker Nkg7 at choroid plexus (CP) in CX3CR1-CreERT2/STOP^flox^-RFP mice. (**e**) Representative confocal images showing immunofluorescence staining of Nkg7 and RFP in untreated control mice and PLX treated mice (PLX^1X^).

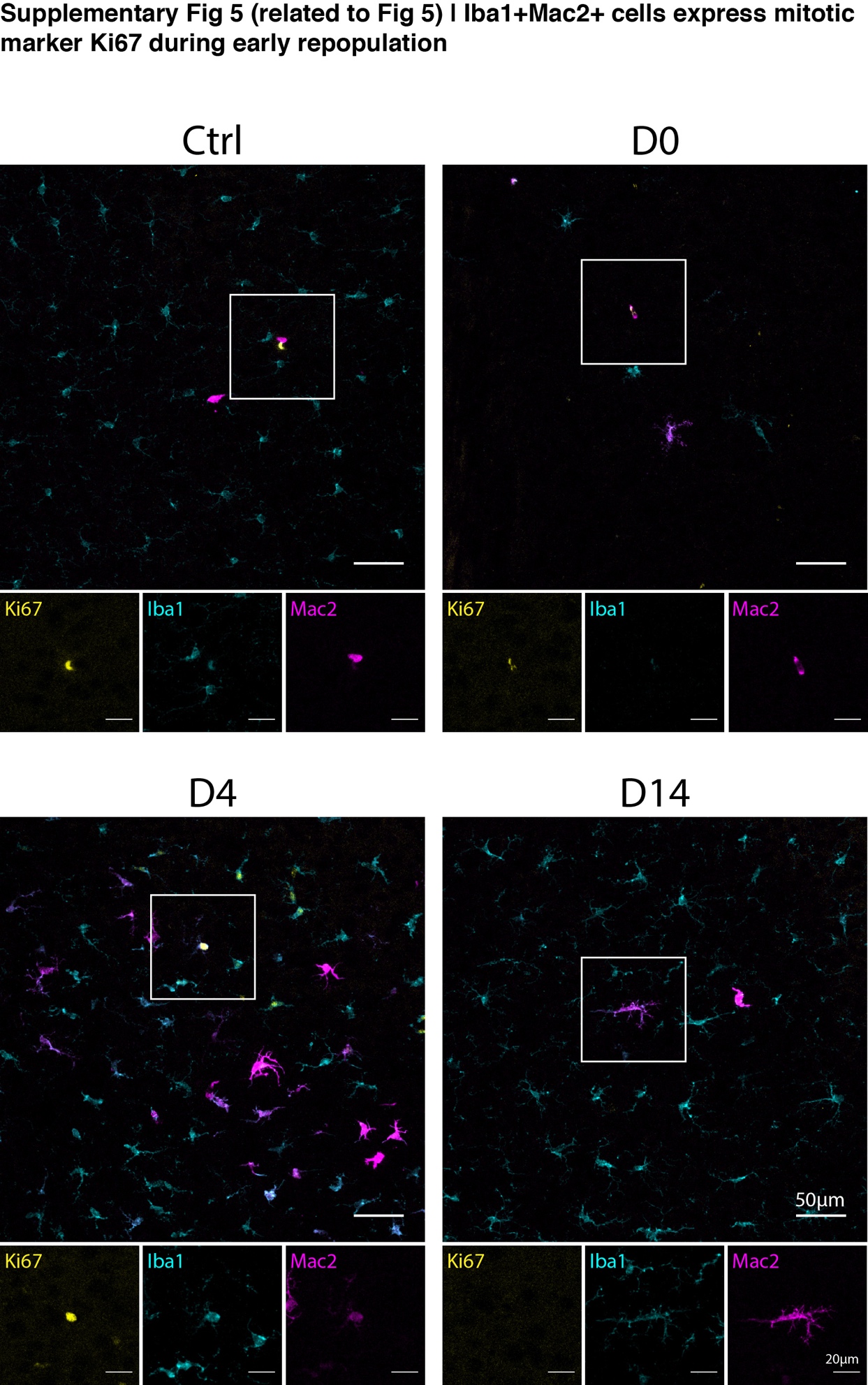

**Supplementary Fig 5 (related to Fig 5) | Iba1+Mac2+ cells express mitotic marker Ki67 during early repopulation**

Representative confocal images showing immunofluorescence staining of Ki-67 (yellow), Iba1 (cyan) and Mac2 (magenta) in naïve C57/BL6J mice, PLX treated mice (D0), mice underwent early microglial repopulation (D4) and longer repopulation (D14). Boxed area is shown by separated channels for each marker at the bottom.

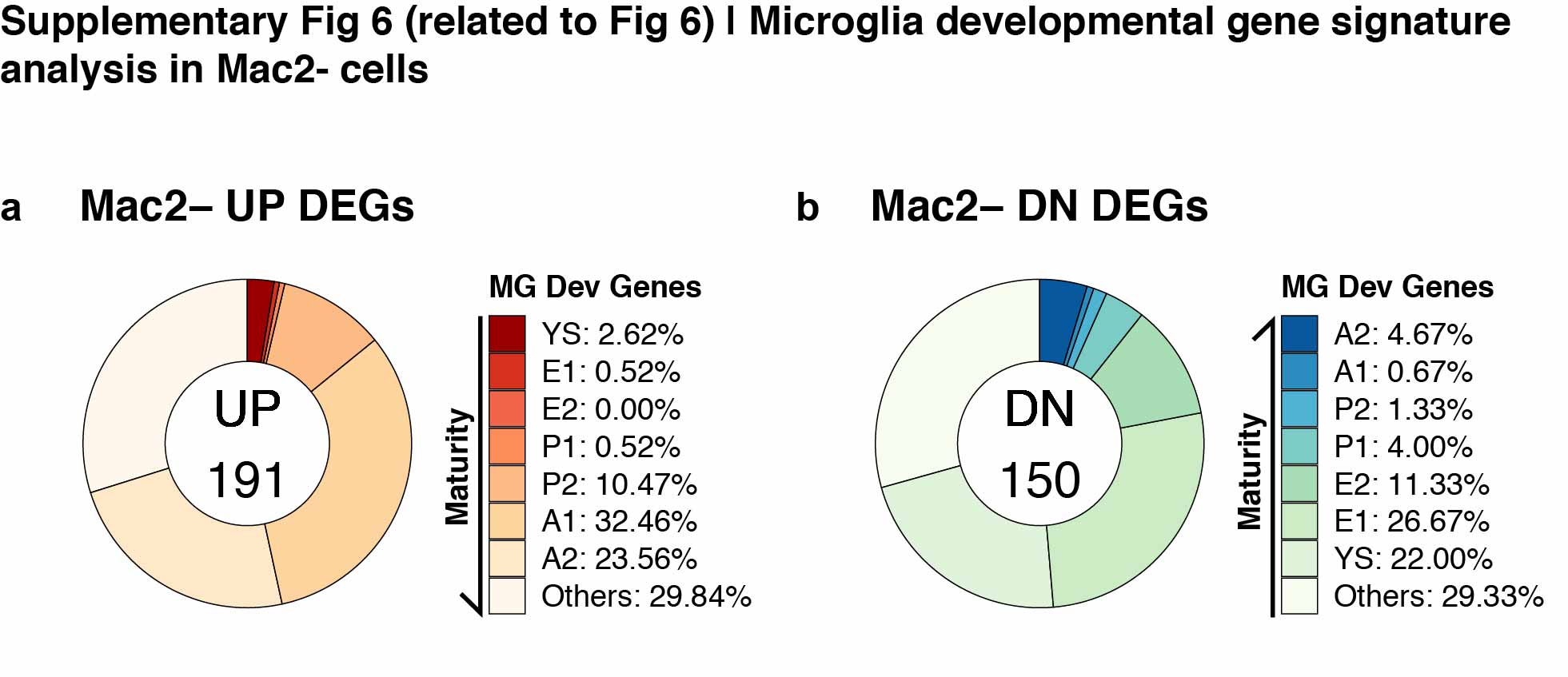

**Supplementary Fig 6 (related to Fig 6) | Microglia developmental gene signature analysis in Mac2- cells**

(**a**) Donut chart showing the percentage of microglial developmental genes in upregulated DEGs found in Mac2- cells (191 genes). (**g**) Donut chart showing the percentage of microglial developmental genes in downregulated DEGs found in Mac2- cells (150 genes).

**Supplementary table 1 | DEG list for all clusters and Mac2+ cells**

DEGs identified in Cluster 1-9, total Mac2+ cells, Mac2+ cell from Ctrl brain
